# Programmed Clonal Expansion Associated with V(D)J Recombination Drives an ATM-Dependent Vulnerability Underlying Preferential Lymphocyte Depletion and Myeloid Bias Following DNA Damage

**DOI:** 10.64898/2026.08.05.743027

**Authors:** Katherine H. Lee, Zhengping Shao, Yunyue Wang, Brian J. Lee, Angelina Li, Faye Yan, Peter A. Sims, Shan Zha

## Abstract

Myeloid bias is a hallmark of aging and genotoxic stress, yet the mechanisms underlying the preferential suppression of lymphopoiesis and the resulting predominance of myeloid cells following DNA damage remain incompletely understood. Here, we systematically characterized the acute hematopoietic response to a clinically relevant 2 Gy dose of ionizing radiation in young adult mice. Forty-eight hours after irradiation, overall bone marrow cellularity was reduced by ∼50–60%, with hematopoietic stem and progenitor cells (HSPCs) and myeloid populations declining proportionally. Strikingly, immature and naïve B cells in the bone marrow and developing T cells in the thymus exhibited profound hypersensitivity, declining by ∼90%, thereby recapitulating the preferential vulnerability of lymphocytes to DNA damage. Single-cell RNA sequencing mapped this vulnerability to cells undergoing programmed clonal expansion associated with V(D)J recombination. Specifically, B and T lymphocytes immediately following productive V(D)J recombination at the immunoglobulin heavy-chain (IgH) and T-cell receptor β (TCRβ) loci were the most radiosensitive, resulting in the marked depletion of the immediately downstream small pre-B and CD4⁺CD8⁺ double-positive (DP) thymocyte populations. Mechanistically, this stage-specific radiosensitivity was mediated by ATM-dependent DNA damage responses, as *Atm* deficiency selectively rescued the hypersensitivity of clonally expanding lymphoid progenitors while leaving the global reduction of HSPCs and myeloid cells largely unchanged. Rapidly proliferating S3 erythroblasts also exhibited a similar ATM-dependent hypersensitivity, suggesting that programmed proliferative bursts may represent a general determinant of radiation sensitivity. Together, these findings identify programmed clonal expansion associated with V(D)J recombination as an intrinsic developmental vulnerability that underlies the preferential suppression of lymphopoiesis following DNA damage and provides a mechanistic explanation for the emergence of myeloid bias.

**Key Points:**

- 2 Gy irradiation preferentially depletes developing mouse B and T cells beyond global hematopoietic loss
- Programmed clonal expansion drives ATM-dependent DNA damage sensitivity in developing lymphocytes.

## Introduction

Over 60% of cancer patients in the United States receive radiation therapy, typically delivered as ∼2 Gy daily fractions^1,2^. Circulating blood cells and hematopoietic stem and progenitor cells (HSPCs) in the bone marrow (BM) inevitably sustain collateral damage. Although peripheral blood counts generally recover within weeks^3^, repeated irradiation imposes selective pressure on HSPCs, promoting clonal hematopoiesis (CH) and expansion of pre-existing clones with leukemogenic potential^4,5^. However, how clinically relevant low-dose radiation acutely affects hematopoietic lineages and developmental stages *in vivo* remains elusive^6^. Meanwhile, the impact of chronic DNA damage on hematopoiesis has been extensively studied during aging^7,8^. A hallmark of aged and chronically damaged HSPCs is myeloid bias, characterized by preferential suppression of lymphopoiesis and relatively increased myeloid output^9–11^. While *in vitro* studies demonstrate preferential myeloid commitment by aged HSPCs using terminal differentiation markers^12,13^, mature peripheral T cells are among the most radioresistant hematopoietic populations^13^, suggesting that radiation hypersensitivity may arise from specific developmental stages or cellular states rather than lymphoid identity itself. Ionizing radiation (IR) induces DNA double-strand breaks (DSB) and oxidative base damage and activates three related protein kinases - ATM, ATR, and DNA-PKcs^14^. While DNA-PKcs functions principally in the non-homologous end-joining DSB repair pathway^15^ and ATR primarily responds to replication stress, ATM phosphorylates chromatin-bound factors (e.g., H2AX, 53BP1), activates cell-cycle, and transcriptional checkpoints to facilitate precise repair of DSBs, including programmed DSBs generated during V(D)J recombination^16–19^. Correspondingly, ataxia-telangiectasia (A-T) patients with germline ATM-deficiency display extreme IR hypersensitivity, frequent immunodeficiency, and lymphoid malignancies^20,21^.

Unlike cultured cells, where IR primarily triggers transient cell-cycle arrest, the hierarchical organization of hematopoiesis translates damage at one developmental stage into downstream depletion and compensatory replenishment. In adult mice, BM hematopoietic stem cells (HSCs) give rise to multipotent progenitors (MPPs), which differentiate into common myeloid progenitors (CMPs), megakaryocyte-erythroid progenitors (MEPs), and common lymphoid progenitors (CLPs), ultimately producing the full spectrum of mature blood cells^22,23^. A defining feature of lymphopoiesis is programmed clonal expansion coupled to V(D)J recombination. In B cells, proliferation occurs before and after productive IgH recombination in early-Pro-B and large-Pre-B cells^24^, whereas in αβ T cells, successful TCRβ recombination similarly triggers clonal expansion before TCRα rearrangement in CD4⁺CD8⁺ double-positive (DP) thymocytes^25^, maximizing antigen receptor diversity with the burst of proliferation.

To understand the mechanism of genotoxic lymphoid suppression, we analyzed young adult mice 48 hr after 2 Gy whole-body irradiation using flow and single-cell RNA-sequencing (scRNA-seq). Beyond the ∼50% depletion observed across most hematopoietic stages and lineages, developing, but not mature lymphocytes exhibited marked, ATM-dependent hypersensitivity to IR during programmed clonal expansion following IgH or TCRβ recombination. Cycling S3 erythroblasts displayed analogous ATM-dependent hypersensitivity, suggesting that proliferation constitutes a shared vulnerability to IR across lineages. Together, these findings reveal programmed clonal expansion during lymphocyte development as a previously unappreciated mechanism contributing to genotoxic lymphoid suppression and the myeloid bias observed after DNA damage and potentially during hematopoiesis aging.

## Methods

### Mice and irradiation

All animal procedures were approved by the Columbia University IACUC. Wild-type (WT) 129S/Sv (129S6/SvEvTac) mice and *Atm^−/−^* mice^26,27^ of the same background were used. Both male and female mice were used and pooled because no sex differences were observed. Mice received whole-body X-ray irradiation (0.8 Gy/min; MultiRad350) and were analyzed 2-8 days later, with primary analyses focused on 2 Gy at 48 h. Mice were housed in a specific pathogen-free facility with ad libitum access to food and water. The animals were randomly assigned to the control and IR groups while keeping the sex balanced, and investigators were not blinded.

### Tissue harvest and peripheral blood analysis

Peripheral blood was collected by submandibular bleeding into EDTA-coated tubes (Microvette 500 K3E) and analyzed on a Genesis hematology analyzer (Oxford Science) for complete blood counts (CBC). Mice were euthanized by CO₂ asphyxiation followed by cervical dislocation. BM was obtained by flushing one femur and one tibia with PBS/1% FBS; total nucleated cellularity was determined using a Countess 3 automated cell counter (Invitrogen). The thymus was weighed at harvest, and the tissue weight was normalized to body weight (tissue weight/body weight × 100). Single-cell thymus suspensions were prepared in PBS/1% FBS and passaged through a 70-µm nylon strainer (ThermoFisher, Cat. 352350). Splenocytes were treated with ACK lysis buffer (Lonza, Cat: BP10-548E) before downstream analyses.

### Flow cytometry

Flow cytometry analyses of hematopoietic cells and lymphocytes were performed using previously published methods with minor modifications^28–31^. BM, spleen, and thymus single-cell suspensions were stained using antibody panels detailed in Supplemental Table 1. For HSPC analysis, BM cells were incubated with a biotin-conjugated lineage depletion cocktail (anti-CD5, CD3ε, CD8α, CD4, B220, Gr-1, Ter119) for 1 hour at 4°C, washed, and then stained with directly conjugated antibodies for 10 minutes at room temperature. All other panels were stained directly without prior lineage depletion and run as separate single-stain experiments; antibodies shared across panels reflect independent staining runs on distinct cell preparations. Erythroid progenitors were staged S0-S5 by CD71 and Ter119 expressions^28,29,32^. BM B cell developmental stages were resolved as: pro-B (B220^+^IgM^−^CD43^+^), pre-B (B220^+^IgM^−^CD43^−^), naïve B (B220^+^IgM^+^), and recirculating B cells (B220^High^IgM^+^). Large and small pre-B cells were distinguished by FSC. Thymic subpopulations were resolved as double negative (DN; CD4^−^CD8^−^), double positive (DP; CD4⁺CD8⁺), CD4⁺ single positive, and CD8⁺ single positive (SP) thymocytes. Data were acquired on an LSR II (BD Biosciences) or Attune NxT (Thermo Fisher) and analyzed with FlowJo v10. Absolute cell counts for flow cytometry-defined populations were calculated by multiplying total tissue cellularity by the frequency of each gated population within the corresponding sample.

### Histology analyses

Tibiae were fixed in 10% neutral buffered formalin for 24 hr, decalcified, and processed for hematoxylin and eosin (H&E) staining by the Molecular Pathology Core Facility at Columbia University Irving Medical Center. Sections were imaged at 200× magnification.

### Cell enrichment for single-cell RNA sequencing

For all scRNA-seq experiments, erythrocytes were depleted by sequential hypotonic lysis (ACK lysing buffer, Lonza, Cat: BP10-548E, 2 min, room temperature) followed by magnetic depletion using anti-Ter119 microbeads (1 µL per 10⁶ cells; Miltenyi Biotec, 130-049-901) on LS columns. All samples contained <3% Ter119⁺ cells at submission. For cKit-enriched BM scRNA-seq, Ter119-depleted BM cells were further processed using anti-CD117 (cKit) microbeads (Miltenyi Biotec, 130-091-224) with a minimum input of 6×10⁶ cells; cKit⁺ cells were retained on column, washed, and eluted. For thymic scRNA-seq, single-cell thymus suspensions were processed by ACK lysis and Ter119 bead depletion only, using the same protocol as BM. Following enrichment, cells were washed three times in Ca²⁺/Mg²⁺-free PBS/10% FBS to remove EDTA, counted, and resuspended at ≥10⁷ cells/mL for immediate transport to the sequencing core. Viability was >70% and input was ≥10⁶ cells per sample. A fraction of each sample was reserved for post-enrichment flow cytometric validation.

### Single-cell RNA sequencing and data processing

Single-cell RNA sequencing was performed at the JP Sulzberger Genome Center using the 10x Genomics Chromium Single Cell 3′ Gene Expression platform. Libraries were sequenced on an Illumina NovaSeq X Plus sequencer. Raw data were demultiplexed and aligned to the mouse reference genome (mm10) using Cell Ranger; gene-by-cell count matrices were loaded into Seurat v5.0.1 in R v4.3.1. Low-quality cells were excluded by the following criteria: >6,000 or <200 genes detected; >50,000 or <500 UMIs; >10% mitochondrial reads. Data were log-normalized, and principal component analysis was performed on the 2,000 most variable genes. The number of PCs used for downstream analysis was selected at the point where cumulative variance exceeded 90% and per-PC contribution fell below 5%, combined with the point where variance difference between consecutive PCs fell below 0.1. Clusters were identified using Seurat FindClusters at resolution 0.8 and visualized by UMAP. Cell types were manually annotated by examining the expression of canonical marker genes across clusters. Marker gene sets for each dataset are provided in Supplemental Table 2, along with the published references from which they were derived. This pipeline was applied to total BM (Ter119-depleted; 2 ctrl, 3 IR WT mice; n=7,691 and n=20,270 cells. cKit-enriched BM (2 ctrl, 2 IR WT mice across two independent experiments; 11,538 ctrl and 11,584 IR cells), and thymus (1 ctrl, 1 IR WT mouse; 8,822 ctrl and 3,142 IR cells). Atm⁻/⁻ total BM scRNA-seq was processed identically (1 ctrl, 1 IR mice Atm⁻/⁻; 7,293 ctrl and 4,054 IR cells).

### Single-cell RNA-seq quantification and statistical analysis

scRNA-seq data were normalized and plotted using the Seurat framework with additional statistical and display tools from other papers^33–35^. Briefly, for scRNA-seq datasets with biological replicates, cell-type abundance was calculated using propeller from the speckle R package with annotated cell types as clusters, sample identifiers as biological replicates, and condition as the experimental group. This analysis was applied to Ter119-depleted total BM and cKit-enriched BM datasets. Cell-type proportions were calculated per sample, and condition-associated differences were evaluated with Benjamini-Hochberg FDR correction across tested populations. For scRNA-seq datasets or comparisons with one biological sample per condition, including thymus and *Atm*^⁻/⁻^ BM, analyses are reported descriptively and interpreted alongside matched flow cytometry data where available. Cell-cycle phase assignment was performed using Seurat CellCycleScoring available at satijalab.org/seurat. Mouse S-phase and G2/M-phase gene sets were obtained from a Mus musculus cell-cycle marker table and mapped from Ensembl gene IDs to gene symbols using AnnotationHub and ensembld. Cells were assigned to G0/G1, S, or G2/M phase based on Seurat scoring, and phase distributions were summarized as the percentage of cells in each phase within each annotated cell type, subcluster, and condition. Differential expressions were performed using Seurat FindMarkers on log-normalized RNA expression values. Genes were ranked by adjusted p value and log₂ fold change. Marker detection frequency was calculated as the percentage of cells within each annotated population with detectable expression of the indicated gene. scRNA-seq analyses and visualizations were performed in R v4.3.1 and figures generated in GraphPad Prism.

### Statistical analysis

Flow cytometry and CBC data were analyzed using unpaired two-tailed Welch’s t-tests. Statistical analyses were performed using GraphPad Prism and R. Data are presented as mean ± SEM unless otherwise indicated. Statistical tests are described in the corresponding figure legends. P < 0.05 was considered statistically significant. Percent change was calculated as [(IR mean − Ctrl mean) / Ctrl mean] × 100.

## Results

### 2 Gy IR causes a ∼50% depletion of nucleated cells and preferential loss of lymphocytes

Pilot studies in young adult mice (8–12 weeks) receiving 1–8 Gy whole-body irradiation identified 2 Gy with 48 h recovery as a reproducible early time point preceding partial hematopoietic recovery by days 4–8 (Fig. S1A-F). Body weight was unchanged at 48 h (Fig. S1G). CBC revealed a modest reduction in circulating red blood cells (RBCs) (23.9%, p=0.03) and hemoglobin (30.0%, p=4.61×10⁻³), accompanied by a small decrease in mean corpuscular hemoglobin (7.6%, p=4.08×10^-4^) and hematocrit (60.7% to 45.2%, p=0.02), consistent with limited hemolysis^36^ (Fig. 1A and S1H-J). Platelet counts were unchanged (Fig. S1K). In contrast, WBCs declined by 61.4% (p=6.03×10⁻³), including marked reductions in short-lived neutrophils (55.8%, 1.65×10^9^ to 7.29×10^8^ cells/L, p=0.02) and long-lived lymphocytes (64.8%, 3.43×10^9^ to 1.21 ×10^9^ cells/L, p=5.25×10⁻³) (Fig. 1B-D). No significant sex-specific differences were detected (Fig. S1L-O), and male and female mice were pooled for subsequent analyses. These findings demonstrate that WBCs are substantially more radiosensitive than RBCs, with lymphocytes exhibiting slightly greater sensitivity than myeloid cells, consistent with previously reported lineage biases^12,37^.

**Figure 1.**
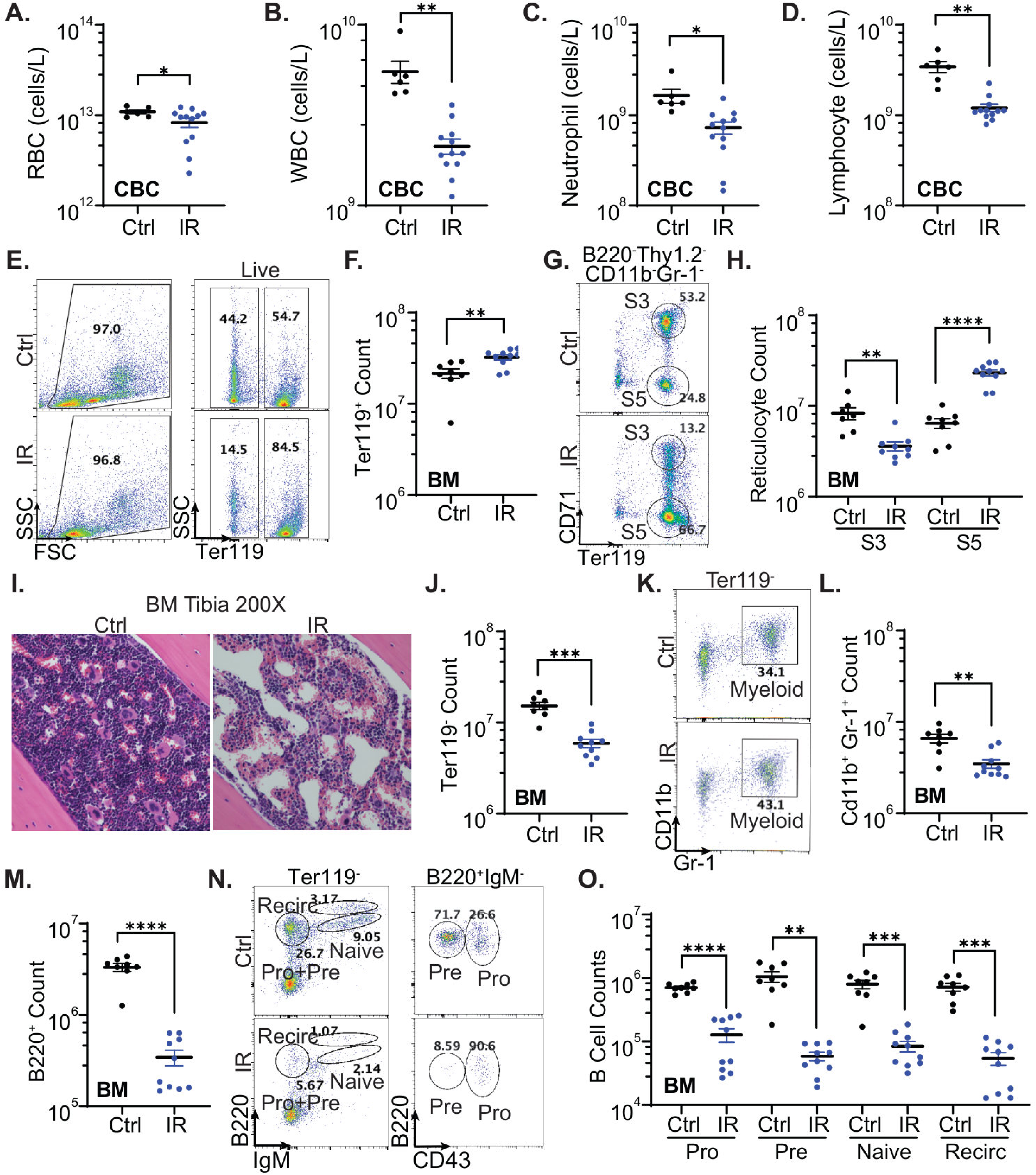
Acute 2 Gy irradiation preferentially depletes peripheral lymphocytes and developing bone marrow B cells. **(A-D)** Peripheral blood complete blood count analysis of red blood cell (RBC), white blood cell (WBC), neutrophil, and lymphocyte counts in control and irradiated mice. **(E)** Representative flow cytometry gating of live bone marrow cells into Ter119⁺ and Ter119⁻ fractions **(G)** and corresponding absolute Ter119⁺ bone marrow counts. **(G)** Representative CD71/Ter119 flow cytometry staging of erythroid cells after exclusion of B220⁺, Thy1.2⁺, CD11b⁺, and Gr-1⁺ cells **(H)** and absolute counts of S3 erythroblasts and S5 erythroid cells. **(I)** H&E-stained tibial bone marrow sections, imaged at 200X magnification. **(J)** Absolute Ter119⁻ bone marrow counts. **(K)** Representative flow cytometry gating of CD11b⁺Gr-1⁺ myeloid cells **(L)** and corresponding absolute counts. **(M)** Absolute B220⁺ bone marrow B cell counts. **(N)** Representative B cell developmental gating of pro-B (B220^+^IgM^−^CD43^+^), pre-B (B220^+^IgM^−^CD43^−^), naïve B (B220^low^IgM^+^), and recirculating B cells (B220^high^IgM^+^) **(O)** and corresponding absolute counts. Data are shown as mean ± SEM. Statistical significance was determined by an unpaired two-tailed Welch’s t-test. ns, not significant; **p < 0.01; ***p < 0.001; ****p < 0.0001.

### IR causes preferential depletion of BM B cell progenitors undergoing clonal expansion

We next analyzed BM, the major site of hematopoiesis, by flow cytometry (n=8 Ctrl and n=10 IR). After IR, the fraction (57.2% to 84.3%) and absolute count (2.15×10^7^ to 3.25×10^7^ cells, p=5.15×10^-3^) of Ter119^+^ erythroid-lineage cells increased (Fig. 1E-F). Murine BM erythropoiesis can be divided into five stages: S0 (uncommitted progenitors, CD71^−^Ter119^−^), S1-S3 (erythroblasts, CD71^+^Ter119^+/-^), and S4-S5 (reticulocytes and enucleated erythrocytes, CD71^mid/-^ Ter119^+^)^32,38^ (Fig. 1G). The increase in Ter119^+^ cells was primarily driven by the relatively mature S5 population, which increased by 260.7% from 6.44×10^6^ to 2.32×10^7^ cells (p=2.06×10^-6^) (Fig. 1H). In contrast, the proliferating S3 erythroblasts decreased by 56.5%, from 8.29×10^6^ to 3.61×10^6^ cells (p=9.07×10^-3^) (Fig. 1H). This pattern is consistent with relative IR resistance among enucleated S5 erythroid cells and increased sensitivity among proliferating erythroblasts^39^. BM histology confirmed mature RBC enrichment and partial hemolysis (200x) (Fig. 1I). Consistent with ∼60% peripheral WBC depletion, Ter119^−^ nucleated BM cells decreased by 61.2% after IR, from 1.52×10^7^ to 5.89×10^6^ cells (p=1.30×10^-4^) (Fig. 1J), also evident by histology (Fig. 1I). Specifically, CD11b^+^Gr-1^+^ myeloid cell counts decreased by 47.3% from 6.60×10^6^ to 3.48×10^6^ cells (p=2.80×10^-3^) (Fig. 1K-L), parallelling the ∼50% reduction in peripheral neutrophils (Fig. 1C) and consistent the short half-life of mature neutrophils^40,41^. Strikingly, B220^+^ B cell counts decreased 89.7% from 3.32×10^6^ to 3.43×10^5^ cells (p=3.00×10^-5^) (Fig. 1M), exceeding the ∼65% decline in peripheral lymphocytes (Fig. 1D). These findings suggest that BM B cells are more IR-sensitive than peripheral B cells.

We next analyzed BM B cell development by FACS. B220^+^ BM B cells can be divided into four major developmental stages based on surface markers from primitive to mature: pro-B (B220^+^IgM^−^CD43^+^), pre-B (B220^+^IgM^−^CD43^−^), naïve B (B220^low^IgM^+^), and recirculating B cells (B220^high^IgM^+^) (Fig. 1N). After IR, all BM B cell compartments were markedly reduced, with the greatest depletion in pre-B cells (-94.4%, 1.05×10^6^ to 5.88×10^4^, p=1.38×10^-3^) followed by naïve B cells (-89.4%), and pro-B cells (-81.9%). IgM+B220^High^ recirculating B cells also decreased 92.4% (7.20×10^5^ to 5.45×10^4^, p=3.15×10^-4^) (Fig. 1O).

### IR causes a proportional ∼50% depletion of HSPC and myeloid compartments

To understand if the ∼50% decline in short-lived myeloid cells reflected depletion at the early progenitors, we analyzed HSPCs using SLAM markers within lineage-negative (Lin⁻) BM cells^31,42,43^ (Fig. 2A). LSK (cKit^+^Sca1^+^) and LK (cKit^+^Sca1^−^) compartments decreased by 64.0% and 65.4% (p<0.01), respectively (Fig. 2A-B). Following IR, the Sca1⁺cKit⁻ (LS) population increased from 1.96×10⁵ to 5.87×10⁵ cells (P=0.05) (Fig. 2A-B). Although the identity and function of these cells remain uncertain, previous studies suggest that this compartment is enriched for lymphoid-biased progenitors and expands in aged mice^44^.

**Figure 2.**
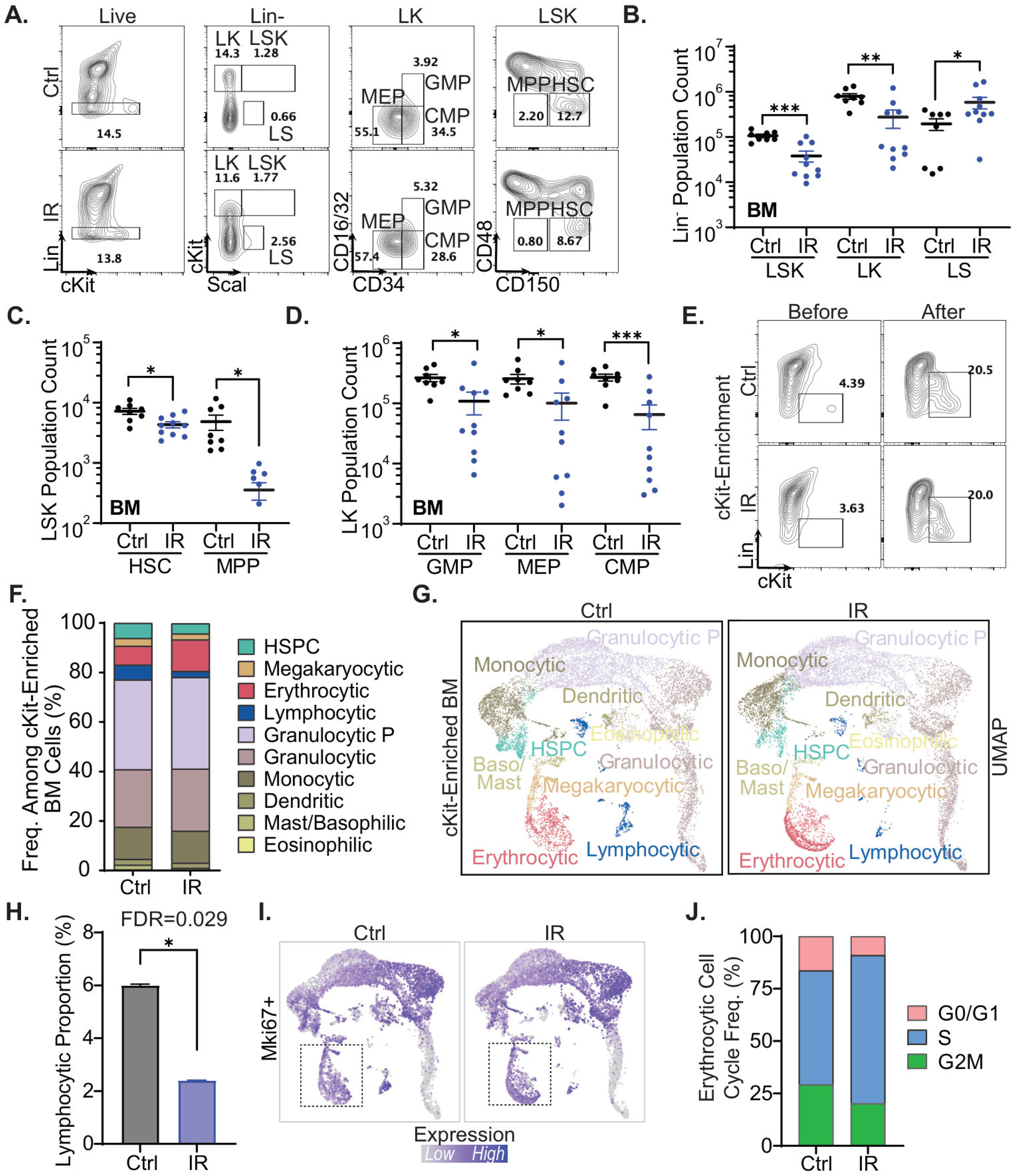
Irradiation preferentially depletes lymphocytic cells in cKit-enriched bone marrow. **(A)** Representative flow cytometry gating of bone marrow lineage-negative (Lin⁻: CD5, CD3ε, CD8α, CD4, B220, Gr-1, Ter119) hematopoietic stem and progenitor cell populations, including LSK, LK, LS, HSC, MPP, GMP, CMP, and MEP compartments. **(B)** Absolute cell counts of Lin⁻ LSK, LK, and LS populations in control and irradiated mice. **(C)** Absolute counts of HSCs and MPPs within the LSK compartment. **(D)** Absolute counts of GMP (CD16/32^+^CD34^+^), CMP (CD16/32^−^CD34^+^), and MEP (CD16/32^−^CD34^−^), within the LK compartment. **(E)** Flow cytometry plots of cKit enrichment before and after magnetic bead enrichment of Ter119-depleted bone marrow cells. **(F-G)** Cell-type frequency and UMAP visualization of cKit-enriched bone marrow scRNA-seq from control and irradiated mice. **(H)** Lymphocytic frequency is significantly reduced in irradiated cKit-enriched bone marrow. **(I)** UMAP visualization of Mki67 expression. **(J)** Cell-cycle phase distribution of erythrocytic cells. Data are shown as mean ± SEM. Statistical significance was determined by unpaired two-tailed Welch’s t-test for flow cytometry data and by replicate-level proportion testing for scRNA-seq cell-type abundance where applicable. *p < 0.05; **p < 0.01; ***p < 0.001.

Within the LSK compartment, less proliferative CD48^−^CD150^+^ HSCs showed a moderate 40.1% decline (7.23×10^3^ to 4.33×10^3^, p=0.01), while the more proliferative CD48^−^CD150^−^ MPPs declined 92.7% (4.85×10^3^ to 3.55×10^2^, p=0.01) (Fig. 2C), consistent with the positive correlation between proliferation and IR sensitivity^45^. Within the more differentiated LK compartment, GMPs (CD16/32^+^CD34^+^), MEPs (CD16/32^−^CD34^−^), and CMPs (CD16/32^−^CD34^+^) decreased by 59.1% (p=0.02), 60.8% (p=0.03), and 75.8% (p=4.38×10^-4^), respectively (Fig. 2D).

ScRNA-seq analysis of cKit-enriched BM cells^46^ resolved 10 major populations and showed largely preserved relative compositions across populations (Fig. 2E-G, Fig. S2A-C). One notable exception was the preferential depletion of the lymphocytic population and mild enrichment of erythroid-lineage cells (Fig. 2F-H). Among erythrocytic cells, the fraction positive for the proliferation marker *Mki67* trended upward from 83.6% to 91.7%, and the S phase fraction increased from 55.6% to 70.6% (Fig. 2I-J, Fig. S2D). Although neither change reached statistical significance, this pattern may reflect feedback upregulation of erythropoiesis. Taken together, FACS and scRNA-seq analyses of BM following IR revealed a proportional ∼50-70% reduction in HSPC and myeloid compartments, contrasting with preferential ∼90% depletion of developing and newly generated B lymphocytes.

### Large pre-B cells undergoing clonal expansion are hypersensitive to IR

To understand the hypersensitivity of developing B cells to IR, we performed scRNA-seq on Ter119-depleted total BM (Ter119^+^ <4%) (Fig. S3A) from two control and three irradiated mice 48 hr after 2 Gy IR, resolving nine major populations from 7,691 Ctrl and 20,270 IR-treated cells (Fig. 3A-B, Fig. S3B). Replicate-level patterns were consistent, so samples were pooled for downstream analyses (Fig. S3C-D). The IR-only sample was omitted in Fig. S3C to avoid visual skewing of the batch overlay due to IR-induced B cell depletion but was retained in all other plots. Consistent with FACS, B lymphocytes (*Cd79a/b, Pax5, Igkc)* were preferentially depleted after IR, decreasing from 21.3% to 3.6% of total BM cells (FDR=3.33×10^-4^). In contrast, the relative frequency of other major populations, including granulocytes (*Ly6g, S100a9*), granulocyte progenitors (*Mpo, Elane*), monocytes (*Cx3cr1*), mast cells (*Fcer1a, Cpa3*), HSPCs (c*Kit, Cd34*), T cells (*Cd3d/g, Trbc2*), and NK cells (*Klrk1, Klre1*), remained largely unchanged after IR, consistent with proportional reductions rather than selective depletion (Fig. 3B, S3E-F).

**Figure 3.**
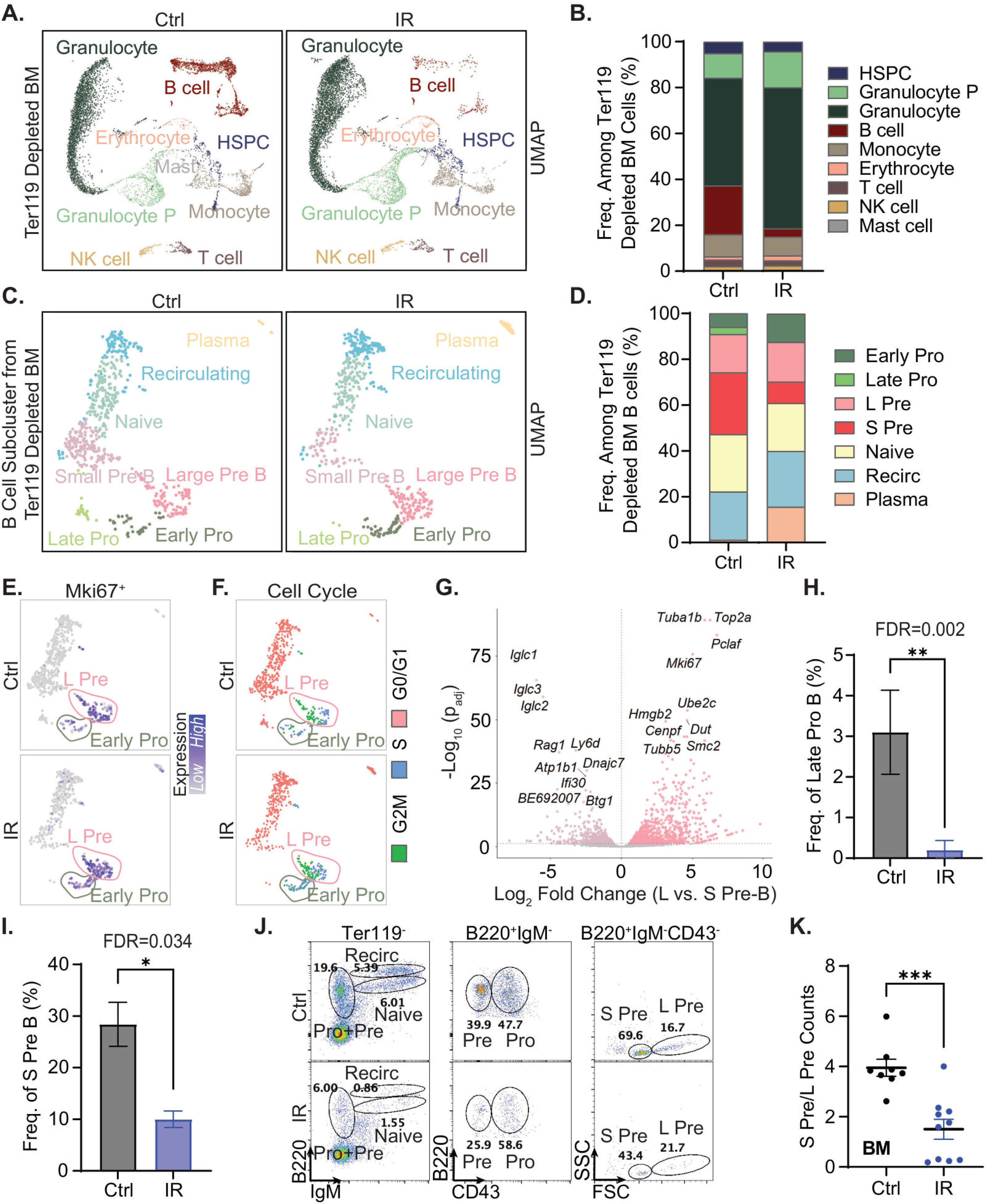
Developing B cells show stage-specific depletion downstream of proliferative expansion after irradiation. **(A)** UMAP visualization of Ter119-depleted bone marrow scRNA-seq from control and irradiated mice, annotated by major hematopoietic populations. **(B)** Relative frequency of annotated Ter119-depleted bone marrow populations. **(C)** UMAP visualization of B cell subclusters from Ter119-depleted bone marrow, annotated across developmental stages from early pro-B cells to plasma cells. **(D)** Relative frequency of annotated B cell populations. **(E-F)** UMAP visualization of Mki67 expression and cell-cycle phase assignment, highlighting proliferative early pro-B and large pre-B populations. **(G)** Percent-detection comparison of genes in large versus small pre-B cells from control bone marrow. Each point represents one gene, and color indicates the difference in detection frequency [large pre-B – small pre-B (% expressing)]. Labeled genes include the representative top 10 enriched and commonly detected markers. **(H-I)** Frequency of control and irradiated late pro-B and small pre-B cells from subclustered B cells. **(J)** Representative flow cytometry gating of bone marrow B cell populations and FSC-based separation of small and large pre-B cells within the B220⁺IgM⁻CD43⁻ pre-B gate. **(K)** Ratio of small pre-B to large pre-B cells counted by flow cytometry. Data are shown as mean ± SEM. Statistical significance was determined by replicate-level proportion testing for scRNA-seq cell-type abundance and by unpaired two-tailed Welch’s t-test for flow cytometry data. *p < 0.05; **p < 0.01; ****p < 0.0001.

To determine which stages of B-cell development were most sensitive to IR, we isolated B cells (*Cd79a/b, Pax5, Igkc*) and resolved developmental states by scRNA-seq. Bone marrow B-cell development alternates between proliferative and recombination-associated G0 phases marked by V(D)J recombination at Igh and light-chain (Igl, predominantly Igk) loci. Early pro-B cells undergo clonal expansion before becoming quiescent late pro-B cells (*Rag1/2⁺, Dntt⁺, Vpreb1⁺, Igh^-/low^*), where Igh recombination occurs. Productive Igh rearrangement triggers another proliferative burst in large pre-B cells (*Mki67⁺, Top2a⁺, Rrm2⁺*), followed by small pre-B cells (*Rag1/2⁺, Igh^high^, Igkc^+/−^*), which undergo Igl recombination. Cells with productive IgH&L rearrangements mature into naïve B cells (*Igkc⁺, Ighm⁺*), while a small population of recirculating B cells (*H2-Aa, Sell, Ms4a4c*) and plasma cells (*Jchain^+^, Igkc^high^, Sdc1*) was also identified (Fig. 3C–E, Fig. S3G).

In BM B cells, the proliferative cell fraction, defined by *Mki67* expression and S phase markers, was highest in the early pro-B and large pre-B compartments, whereas the subsequent late pro-B and small pre-B stages were largely quiescent and V(D)J recombination-associated (Fig. 3E-F). In controls, early pro-B and large pre-B cells were largely assigned to S and G2/M phases and showed nearly universal expression of proliferative markers, including *Rrm2* (93.0% in pooled early pro-B and large pre-B vs. 3.6% in other B cell subtypes), *Mki67* (98.7% vs. 6.1%), and *Top2a* (97.6% vs. 5.4%). In contrast, late pro-B and small pre-B cells were predominantly assigned to G0/G1 in controls (94.8% and 97.1%) and showed preferential expression of recombination-associated genes, including *Rag1* (72.8% in pooled late pro-B and small pre-B vs. 9.1% in other B cell subtypes) and *Rag2* (33.1% vs. 13.2%) (Fig. 3E-F). Differential expression analysis showed distinct biological programs between large and small pre-B cells, with proliferation and DNA replication genes enriched in large pre-B cells, and recombination-associated and light-chain genes, including *Rag1/2* and *Iglc1/2/3*, enriched in small pre-B cells (Fig. 3G).

At 48 hr after IR, depletion was greatest in the developmental stages immediately following proliferative clonal expansion, namely late pro-B and small pre-B cells marked by *Rag1/2* expression and G0/G1 enrichment (Fig. 3G-I, Fig. S3H). Naïve B cell populations were also reduced, but the degree of depletion declined with developmental distance from the large pre-B stage, with greater loss in naïve B cells than in recirculating or plasma cells (Fig. 3C-D). Because small pre-B cells are typically smaller than cycling large pre-B cells, FACS-based size analysis within the B220^+^IgM^−^CD43^−^ pre-B gate confirmed preferential loss of small pre-B cells within the pre-B compartment: the small pre-B/large pre-B ratio decreased from 3.95 in controls to 0.34 after IR (p=2.472×10^-4^) (Fig. 3J-K). Together, these results suggest that among the BM developing B cells, early pro-B cells and large pre-B cells undergoing clonal expansion are particularly sensitive to IR.

### Thymic T-cell development shows analogous stage-specific depletion after IR

We next asked whether clonal expansion-associated radiosensitivity extends to T cell development, where thymic maturation from double negative (DN) to double positive (DP) and single positive (SP) thymocytes provides an analogous hierarchy. At 48 hr after IR, total thymic cellularity decreased by 60.8% (p=9.59×10^-5^) (Fig. 4A), with a corresponding reduction in thymus/body weight (Fig. S4A). Flow cytometry distinguished DN (*CD4⁻CD8⁻*), DP (*CD4⁺CD8⁺*), CD4⁺ SP, and CD8⁺ SP thymocytes (Fig. 4B-D). Among these populations, DP thymocytes, which arise after clonal expansion following successful TCRβ rearrangement, were the most depleted, decreasing by 78.1% (p=3.39×10^-5^) (Fig. 4D). In contrast, DN, CD4⁺ SP, and CD8⁺ SP thymocytes were not significantly changed (Fig. 4D). The more stage-restricted impact may reflect thymocyte positive and negative selection dynamics that prolong residence within developmental states.

**Figure 4.**
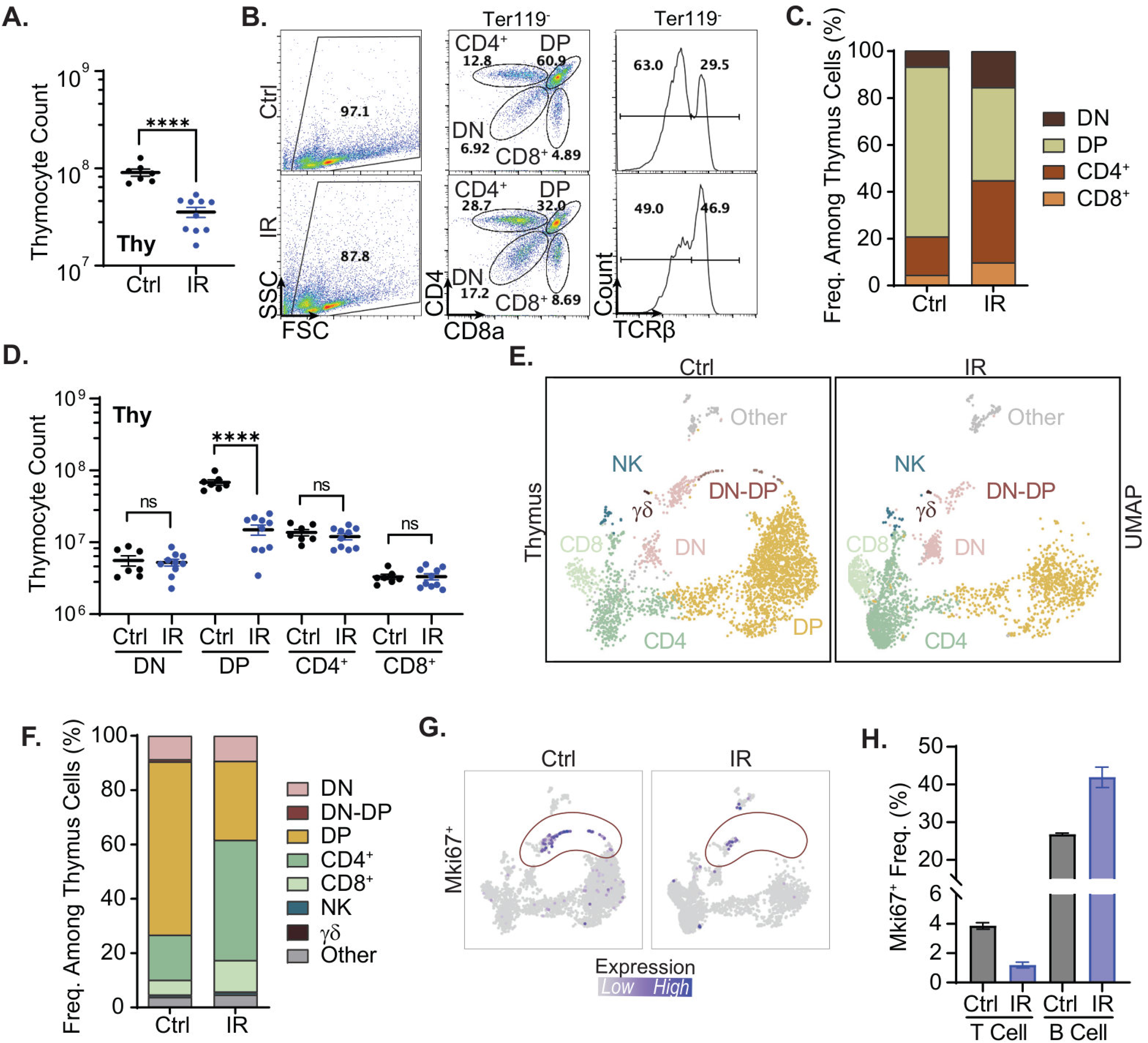
DP thymocytes are selectively reduced 48 hr after irradiation. **(A)** Total thymocyte counts in control and irradiated mice. **(B-C)** Representative flow cytometry gating and relative frequency of DN, DP, CD4⁺ SP, and CD8⁺ SP thymocyte populations. **(D)** Absolute counts of DN, DP, CD4⁺ SP, and CD8⁺ SP thymocytes. **(E-F)** UMAP visualization and relative frequency of DN (*Spi1*), DN-DP (*Mki67, Top2a, Rrm2*), DP (*Cd4, Cd8a, Rag1, Rag2*), CD4⁺ (*Cd4, Cxcr5, Tox, Ctla4*), CD8⁺ (*Cd8a*), and NK cells (*Klrk1, Ncr1*). **(G)** UMAP visualization of Mki67 expression. **(H)** Frequency of Mki67⁺ cells among thymocytes and bone marrow B cells, shown for comparison. Data are shown as mean ± SEM. Statistical significance was determined by an unpaired two-tailed Welch’s t-test. ns, not significant; **p < 0.01; ****p < 0.0001.

Next, we performed thymus scRNA-seq using one control mouse (8,822 cells) and one IR-treated mouse (3,142 cells). We identified six populations: DN (*Spi1*), DN-DP (*Mki67, Top2a, Rrm2*), DP (*Cd4, Cd8a, Rag1, Rag2*), CD4⁺ (*Cd4, Cxcr5, Tox, Ctla4*), CD8⁺ (*Cd8a*), and NK cells (*Klrk1, Ncr1*) (Fig. 4E-F, Fig. S4B). Two clusters exhibited mixed non-T cell identities, co-expressing pDC-associated markers (*Siglech, Cst3, Irf8, Tcf4*), myeloid marker *Spi1*, antigen-presenting cell marker *H2-Aa*, and B cell-associated marker *Iglc3* (Fig. S4C). These clusters represented <5% of cells across conditions (Fig. S4D), were labelled “Other,” and were excluded from subsequent analyses. Consistent with flow cytometry data, DP thymocytes showed the largest proportional decrease after IR, from 63.8% to 29.1%, whereas CD4⁺ SP and CD8⁺ SP fractions increased from 16.6% to 44.2% and 5.4% to 11.6%, respectively (Fig. 4F). DN cells remained similar between conditions (8.7% to 9.3%), as did NK cells (0.8% to 0.9%). As observed in the B cell sub-clustering analysis, the apparent increase in the SP fraction is best interpreted as compositional redistribution rather than expansion, since total thymic cellularity decreased significantly and absolute SP counts were preserved rather than increased.

Given the known proliferative burst after successful TCRβ rearrangement, we separately analyzed the *Mki67*⁺ population, which localized at the DN-DP interface on the UMAP (Fig. 4G). Consistent with the slower dynamics of immature T cell development, the fraction of *Mki67*⁺ thymocytes was smaller than that observed among developing BM B cells. This proliferating population was the most depleted at 48 hr after IR, decreasing from 3.4% to 0.8% of thymocytes, in contrast to the largely replenished proliferative compartment observed in B cells at the same time point (Fig. 4G-H, Fig. 3E-F). These results indicate that clonal expansion during T cell development also confers heightened IR sensitivity, similar to the pattern observed in B cells.

### ATM deficiency selectively attenuates the preferential ablation of developing B cells after IR

To determine whether the hypersensitivity of developing lymphocytes to IR is mechanistically distinct from the ∼50% reduction observed in HSPCs and myeloid cells (Fig. 1), we tested whether ATM kinase, a master regulator of the IR-induced DNA damage response and G2/M checkpoint^47^, partially rescues lymphocyte depletion. Because *Atm^−/−^* mice routinely develop lethal lymphomas by 3 months of age^27,48^, we compared 5-week-old *Atm^+/+^* and *Atm^−/−^* littermates. Consistent with lymphocytopenia of *Atm^−/−^* mice and A-T patients^27,49,50^, untreated *Atm^⁻/⁻^* mice exhibited fewer circulating WBCs, primarily due to reduced lymphocytes (3.01×10⁹ vs. 6.81×10⁹/L; p=0.01), whereas RBC counts, hemoglobin, and neutrophils were comparable (Fig. S5A-C). Following IR, depletion of total WBCs and lymphocytes was significantly attenuated in *Atm^⁻/⁻^* mice, whereas neutrophil depletion was unaffected (Fig. S5A-C).

Next, we analyzed BM by flow cytometry. Overall, nucleated Ter119^−^ BM cells decreased in both *Atm^−/−^* (-61.5%) and *Atm^+/+^* (-74.6%) mice (p=0.90) (Fig. 5A; Fig. S5F). CD11b^+^Gr-1^+^ myeloid cells (-56.9% in *Atm^−/−^* vs. -47.7% in *Atm^+/+^* mice; p=0.52) and HSPCs, including LK cells (-82.7% in *Atm^−/−^* vs. -78.2% in *Atm^+/+^* mice; p=0.55) and LSK cells (-83.7% in *Atm^−/−^* vs. -79.3% in *Atm^+/+^* mice; p=0.35), were similarly affected regardless of *Atm* genotype (Fig. 5B; Fig. S5D-F). In contrast, ATM deficiency significantly attenuated the preferential loss of immature B cells after IR, despite lower baseline B220^+^ B cell numbers in *Atm^−/−^* mice compared with *Atm^+/+^* mice, consistent with lymphocytopenia (p=1.94×10^-3^) (Fig. 5C-D). Specifically, B220^+^ cells were depleted by 90.5% in *Atm^+/+^* but only 70.3% in *Atm^−/−^* mice, leaving significantly more residual BM B220^+^ cells after IR (1.26×10^6^ vs. 8.10×10^5^ in *Atm^+/+^*; p=5.19×10^-3^) (Fig. 5D). Most notably, ATM deficiency preserved small pre-B cells and maintained the small/large pre-B ratio after IR (Fig. 5E; Fig. S5G-H). ScRNA-seq analyses of *Atm^−/−^* BM after IR also confirmed selective rescue of the IR-sensitive late-pro-B population, which decreased from 3.5% to 0.3% in *Atm^+/+^* vs. 7.0% to 4.0% in *Atm*^−/−^ mice, and the small pre-B population, which decreased from 28.4% to 10.0% in *Atm^+/+^* vs. 27.7% to 25.3% in *Atm*^−/−^ mice (Fig. 5F-G; Fig. S5I). This attenuation was not due to reduced baseline proliferation, as *Mki67*^+^ and S-phase fractions in early pro-B and large pre-B cells were similar between untreated *Atm^+/+^* and *Atm^−/−^* mice (Fig. S5J-K).

**Figure 5.**
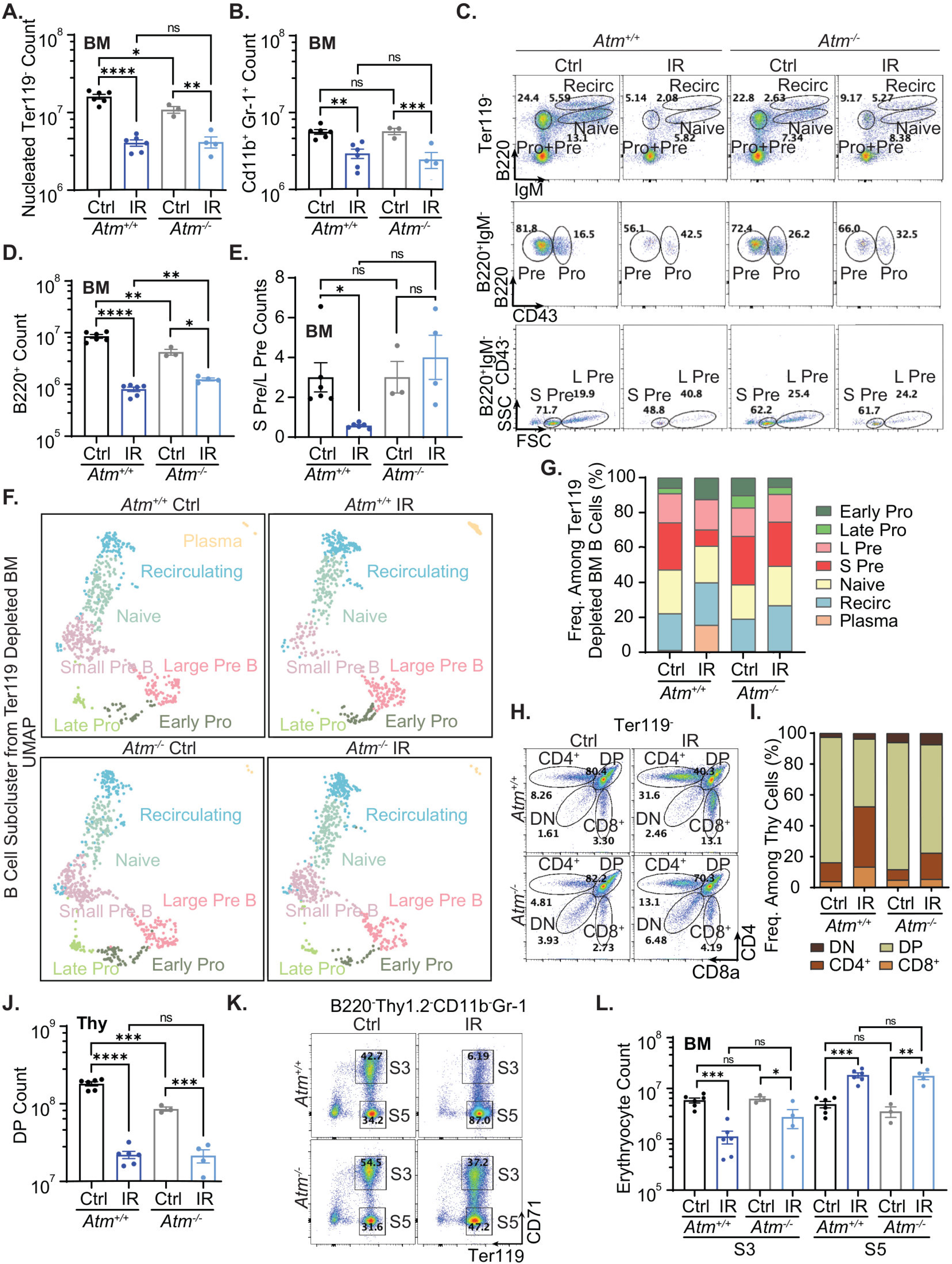
ATM deficiency attenuates preferential attrition of developing lymphocytes and S3 erythroblasts after irradiation. **(A-B)** Absolute counts of bone marrow nucleated Ter119^−^ bone marrow cells and CD11b^+^Gr-1^+^ myeloid cells. **(C)** Representative bone marrow B cell developmental gating. **(D)** Absolute B220^+^ bone marrow B cell counts. **(E)** Ratio of small pre-B to large pre-B cell bone marrow counts. **(F-G)** UMAP visualization and relative frequency of B cell developmental subclusters from Ter119-depleted bone marrow scRNA-seq in *Atm^+/+^* and *Atm^−/−^* mice. **(H-I)** Representative thymus flow cytometry gating and relative frequency of DN, DP, CD4^+^ SP, and CD8^+^ SP thymocytes in *Atm^+/+^*and *Atm^−/−^* mice. **(J)** Absolute DP thymocyte counts. **(K)** Representative CD71/Ter119 erythroid staging in flow cytometry. **(L)** Absolute S3 erythroblast and S5 erythroid cells in *Atm^+/+^* and *Atm^−/−^* bone marrow. Data are shown as mean ± SEM. Statistical significance was determined by an unpaired two-tailed Welch’s t-test. ns, not significant; *p < 0.05; **p < 0.01; ***p < 0.001; ****p < 0.0001.

Consistent with V(D)J recombination defects^27,48,49^, untreated *Atm^−/−^* mice had 53.2% fewer *Thy1.2^+^* thymocytes than *Atm^+/+^* mice, driven by ∼50% decrease of both DP and SP T cells (Fig. 5H-J; Fig. S5L-M). Despite this baseline defect, *Atm^−/−^* mice retained similar DP thymocyte numbers after IR, reflecting reduced proportional depletion in *Atm^−/−^* mice (-75.0% vs. -87.8% in *Atm^+/+^*) (Fig. 5I). The rescue of rapidly proliferating populations was not limited to lymphocytes. IR induced S3 depletion was also attenuated in *Atm^−/−^*(-56.2%) compared with *Atm^+/+^* (-80.7%) mice (Fig. 5K-L), consistent with an ATM-dependent attenuation of proliferating S3 erythroblast depletion. Meanwhile, *Atm*-deficiency did not affect the IR-induced increase in S5 (Fig. 5K-L). Together, these data suggest that an ATM-dependent DNA damage response underlies the preferential depletion of developing lymphocytes, naïve lymphocytes, and rapidly expanding erythroblasts following irradiation.

## Discussion

Our study identifies two distinct components of the hematopoietic response to irradiation: an ATM-independent ∼50–60% reduction across virtually all hematopoietic compartments regardless of proliferation, likely reflecting a general apoptotic response (Fig. 1-2), and an ATM-dependent hypersensitivity of rapidly proliferating lymphoid/erythroid progenitors, which accounts for the myeloid bias and greater sensitivity of developing lymphoid progenitors compared with mature lymphocytes. Specifically, populations immediately following clonal expansion, including late pro-B, small pre-B after IgH rearrangement, and CD4⁺CD8⁺ DP thymocytes after TCRβ rearrangement, were the most severely depleted (Fig.3-4), identifying programmed clonal expansion between V(D)J recombination as a previously unrecognized determinant of radiation sensitivity during lymphocyte development and the resulting myeloid bias.

Mechanistically, ATM kinase drives the preferential depletion of proliferating compartments, as *Atm*-deficiency selectively attenuated radiosensitivity in clonally expanding B- and T-cell progenitors without rescuing the broader 50-60% hematopoietic reduction (Fig.5). Similar protection was observed in rapidly proliferating S3 erythroblasts^51^, suggesting that transient bursts of proliferation represent a shared mechanism of vulnerability across lineages. However, proliferation alone is unlikely to account for all the lineage differences, as myeloid progenitors are also proliferative but at a slower, steadier rate and lack the abrupt transitions between G0/G1 arrest with stable RAG expression and rapid clonal expansion that characterize developing lymphocytes^52^. We speculate that rapid cell-cycle re-entry following G0/G1 arrest imposes an unusually high requirement for genome integrity, rendering expanding clones particularly susceptible to genotoxic stress.

The protective effect of *Atm*-deficiency may appear paradoxical given the profound radiation hypersensitivity of patients with ataxia-telangiectasia and *Atm^−/−^* mice^47,50,53,54^. Our analysis focused on the early hematopoietic response, when ATM-dependent checkpoint effects predominate over long-term survival. Similarly, Trp53- and p21-deficient hematopoietic stem cells exhibit transient protection after genotoxic stress despite impaired long-term fitness^55,56^. Moreover, early (2-day) mortality in irradiated *Atm^⁻/⁻^* mice primarily reflects gastrointestinal toxicity, whereas hematopoietic failure develops later because enucleated mature erythrocytes (T_1/2_=40 day) and platelets are relatively radioresistant^50,57^. Radiation-induced hematopoietic failure is usually most pronounced at approximately 2 weeks after initial exposure. Together, the timing and hematopoiesis focus explain why we primarily observed a protective effect of *Atm*-loss, rather than hypersensitivity.

Although lymphocyte development is well-defined by flow cytometry, scRNA-seq distinguished proliferating early pro-B and large pre-B cells from quiescent, recombination-associated late pro-B and small pre-B cells, captured rare *Mki67⁺* DN-to-DP transitional thymocytes, and identified stress erythropoiesis feedback responses not captured by flow. Quantification of cycling fractions further suggested that developing B cells are more radiosensitive than thymocytes because BM B cells have a larger proliferating compartment than thymocytes, whose development is further shaped by positive and negative selection^58^. Several limitations merit acknowledgment. Megakaryocyte progenitors were largely absent because their large size and irregular morphology limit capture by both scRNA-seq and conventional flow cytometry. In addition, scRNA-seq cannot detect the rapid ATM-dependent post-translational signaling that initiates the DNA damage response, capturing only part of its downstream transcriptional consequences.

Together, our findings establish programmed clonal expansion coupled with V(D)J recombination as a lineage-specific vulnerability to radiation and provide a mechanistic explanation for DNA damage-induced selective ablation of “developing” lymphocytes and the resulting myeloid bias. These results raise the possibility that transient cell-cycle arrest could protect lymphoid progenitors and erythropoiesis during genotoxic therapy. Indeed, a recent clinical report demonstrated that CDK4/6 inhibitors significantly mitigate chemotherapy-induced expansion of TP53-deficient clones in patients^59^.

## Supporting information

Sup Infomration Table 2

## Acknowledgments

We thank Drs. Lei Ding, Lorraine Symington, and Matthew J. Yousefzadeh for insightful suggestions on the thesis committee, and members in the Zha lab for discussion and input. We also thank Dr. Emmanuel Passegué and the Columbia Stem Cell Core for access to and use of the CBC analyzer. This work was partly supported by NIH R01 CA226852, CA293675, CA271595, CA275184, and 1P01CA174653 to SZ, and U54CA274506 to PS. SZ was a scholar of the Leukemia & Lymphoma Society (LLS). K.L was supported by NIH-T32 #1T32CA265828-01A1. This research used the Genomic core at the shared resources funded through the NIH/NCI Cancer Center Support Grant P30CA013696 to the HICCC of Columbia University.

## Authorship Contributions

K.L. and S.Z. designed the experiments, and K.L. analyzed almost all the *Atm*-proficient and deficient murine models and prepared the samples for scRNA-seq analyses. Z.S. established and tested the initial IR dosing and conditions for scRNA-seq in *Atm*-deficient mice. Y.W. analyzed the initial scRNA-seq data. B.J.L., A.L., and F.Y. assisted with colony management, and B.J.L. helped with data management and deposition. P.A.S. advised on scRNA-seq analyses. K.L. and S.Z. wrote the manuscript.

## Supplemental Figure Legends

**Supplemental Figure 1.**
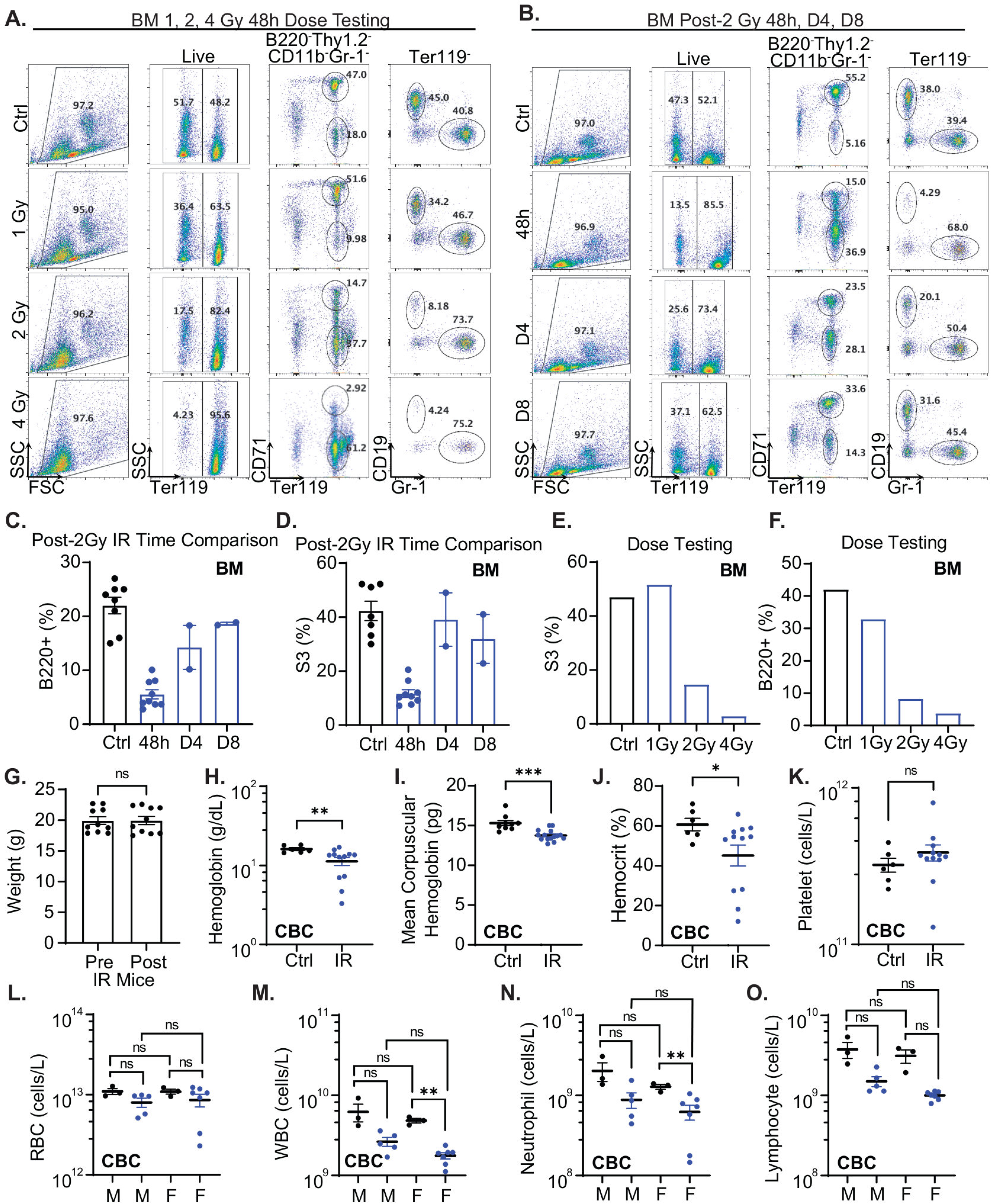
Dose, time course, body weight, and sex-stratified analysis of the acute hematopoietic response to irradiation. **(A)** Representative bone marrow flow cytometry plots 48 hr after 1, 2, or 4 Gy whole-body irradiation. **(B)** Representative bone marrow flow cytometry plots at 48 hr, 4 days, and 8 days after 2 Gy irradiation. **(C-D)** Time course analysis of B220⁺ bone marrow B cell frequency and S3 erythroblast frequency after 2 Gy irradiation. **(E-F)** Dose-response analysis of S3 erythroblast frequency and B220⁺ bone marrow B cell frequency 48 hr after irradiation. **(G)** Body weight before and 48 hr after irradiation. **(H-K)** Peripheral blood parameters in control and irradiated mice, including hemoglobin, mean corpuscular hemoglobin, hematocrit, and platelet count. **(L-O)** Sex-stratified complete blood count analysis of RBCs, WBCs, neutrophils, and lymphocytes in control and irradiated mice. Data are shown as mean ± SEM. Statistical significance was determined by an unpaired two-tailed Welch’s t-test. ns, not significant; *p < 0.05; **p < 0.01; ***p < 0.001.

**Supplemental Figure 2.**
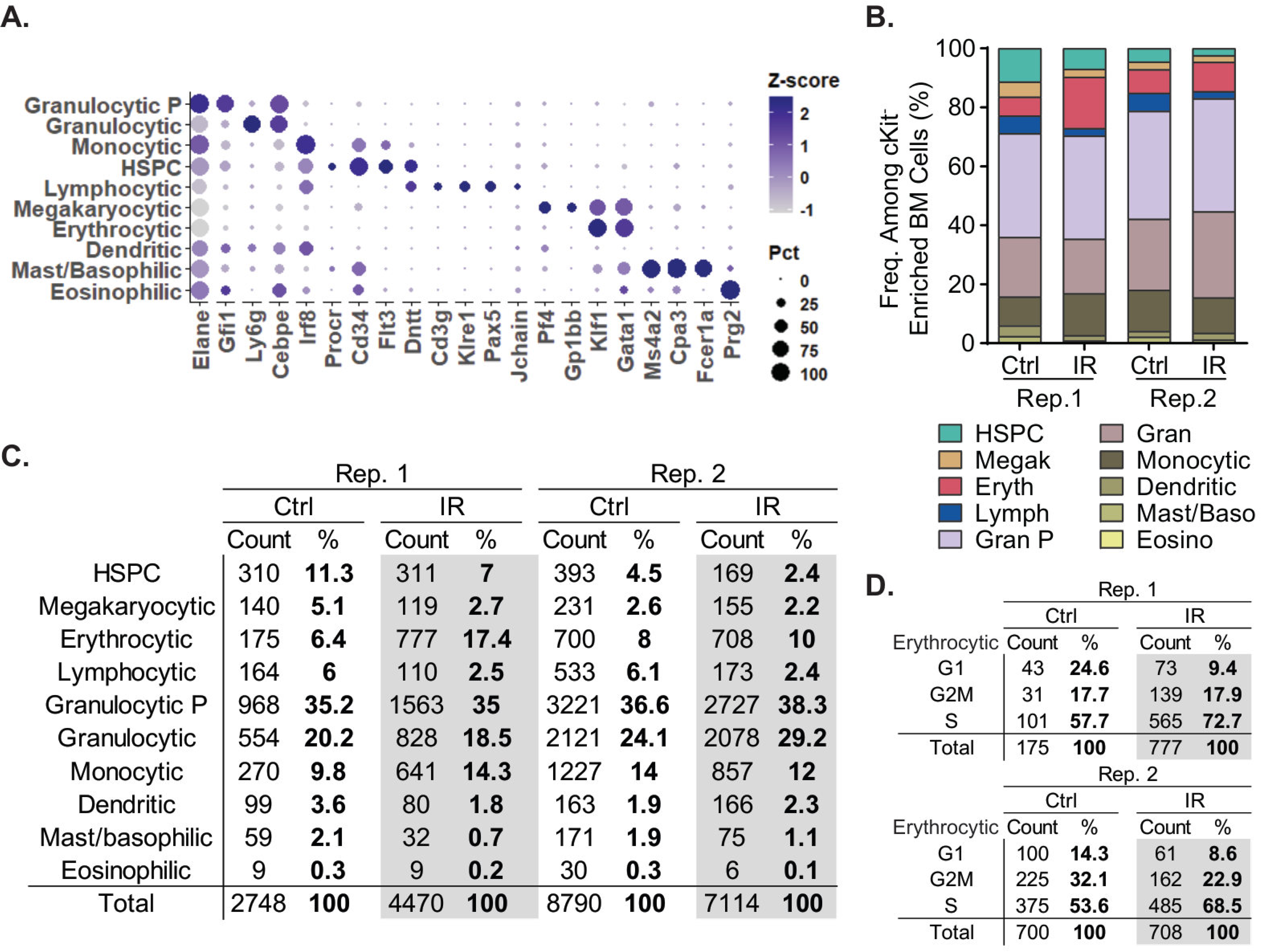
Marker-based annotation and replicate-level composition of cKit-enriched bone marrow scRNA-seq. **(A)** Dot plot showing canonical marker gene expression used to annotate major populations in cKit-enriched bone marrow scRNA-seq. Dot size indicates the percentage of cells expressing each gene, and color indicates scaled expression. **(B)** Replicate-level cell counts and relative frequencies for each annotated population in control and irradiated cKit-enriched bone marrow samples. **(C)** Replicate-level relative frequencies and corresponding cell counts for annotated major hematopoietic populations. **(D)** Replicate-level relative frequencies and corresponding cell counts for erythrocytic cell cycle phases.

**Supplemental Figure 3.**
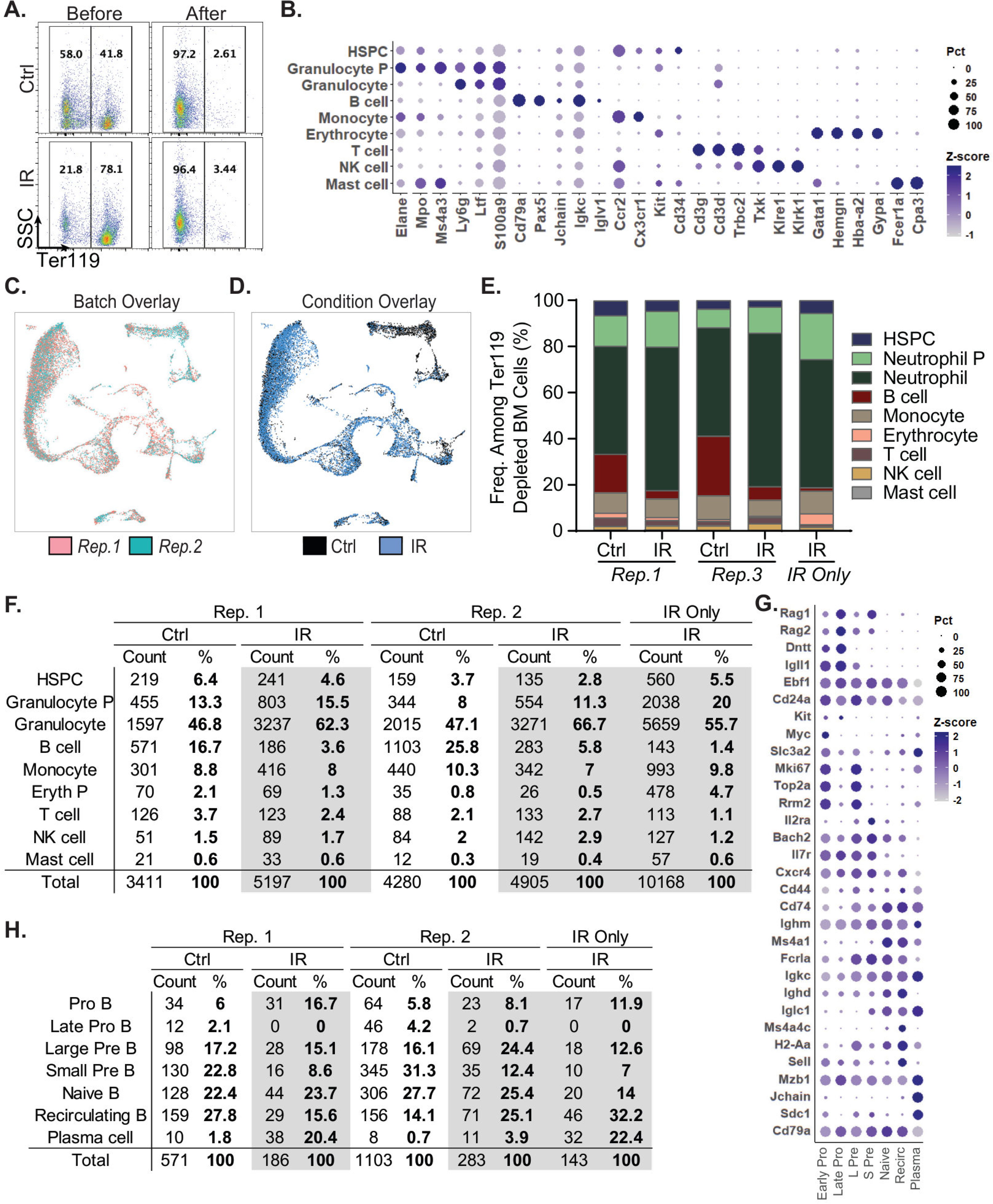
Quality control, annotation, and replicate-level composition of Ter119-depleted bone marrow scRNA-seq and B-cell sub clustering. **(A)** Flow cytometric validation of Ter119 depletion before and after enrichment in control and irradiated bone marrow samples. **(B)** Dot plot shows canonical marker gene expression used to annotate major hematopoietic populations in Ter119-depleted bone marrow scRNA-seq. Dot size indicates the percentage of cells expressing each gene, and color indicates scaled expression. **(C-D)** UMAP overlays showing replicate identity and experimental condition across Ter119-depleted bone marrow scRNA-seq samples. The IR-only sample was omitted in C to avoid visual skewing of the batch overlay due to IR-induced B cell depletion but was retained in all other plots. **(E-F)** Replicate-level relative frequencies and corresponding cell counts for annotated major hematopoietic populations. **(G)** Dot plot showing canonical marker gene expression used to annotate B-cell developmental subclusters. **(H)** Replicate-level cell counts and relative frequencies for B-cell developmental populations in control and irradiated samples.

**Supplemental Figure 4.**
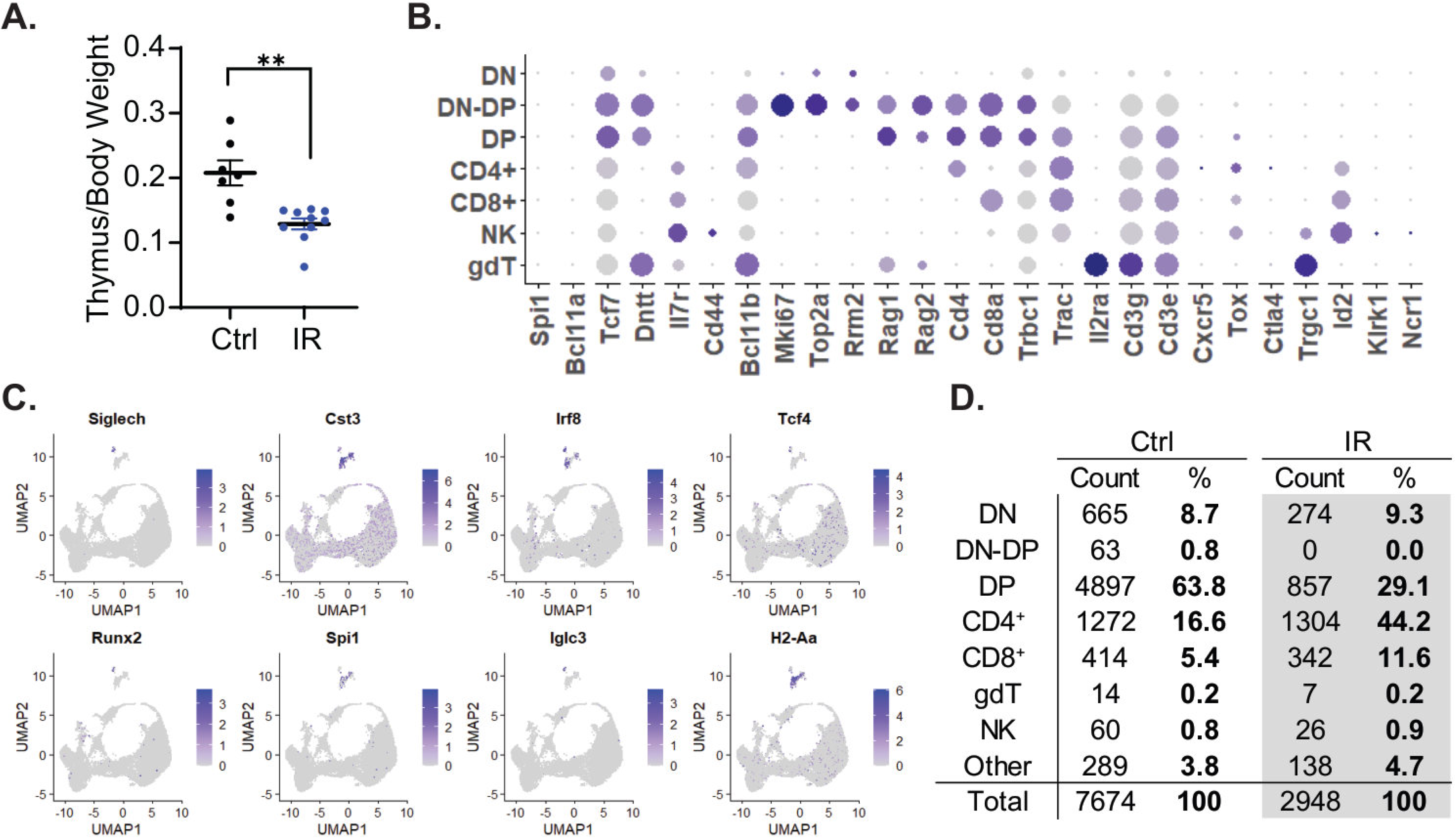
Thymic weight and scRNA-seq annotation after irradiation. **(A)** Thymus-to-body weight ratio in control and irradiated mice. **(B)** Dot plot showing canonical marker gene expression used to annotate thymic populations in scRNA-seq. Dot size indicates the percentage of cells expressing each gene, and color indicates scaled expression. **(C)** Feature plots showing expression of marker genes used to identify the “Other” cluster, including pDC-associated, myeloid, antigen-presenting, and B cell-associated genes. **(D)** Cell counts and relative frequencies of annotated populations in control and irradiated thymus scRNA-seq.

**Supplemental Figure 5.**
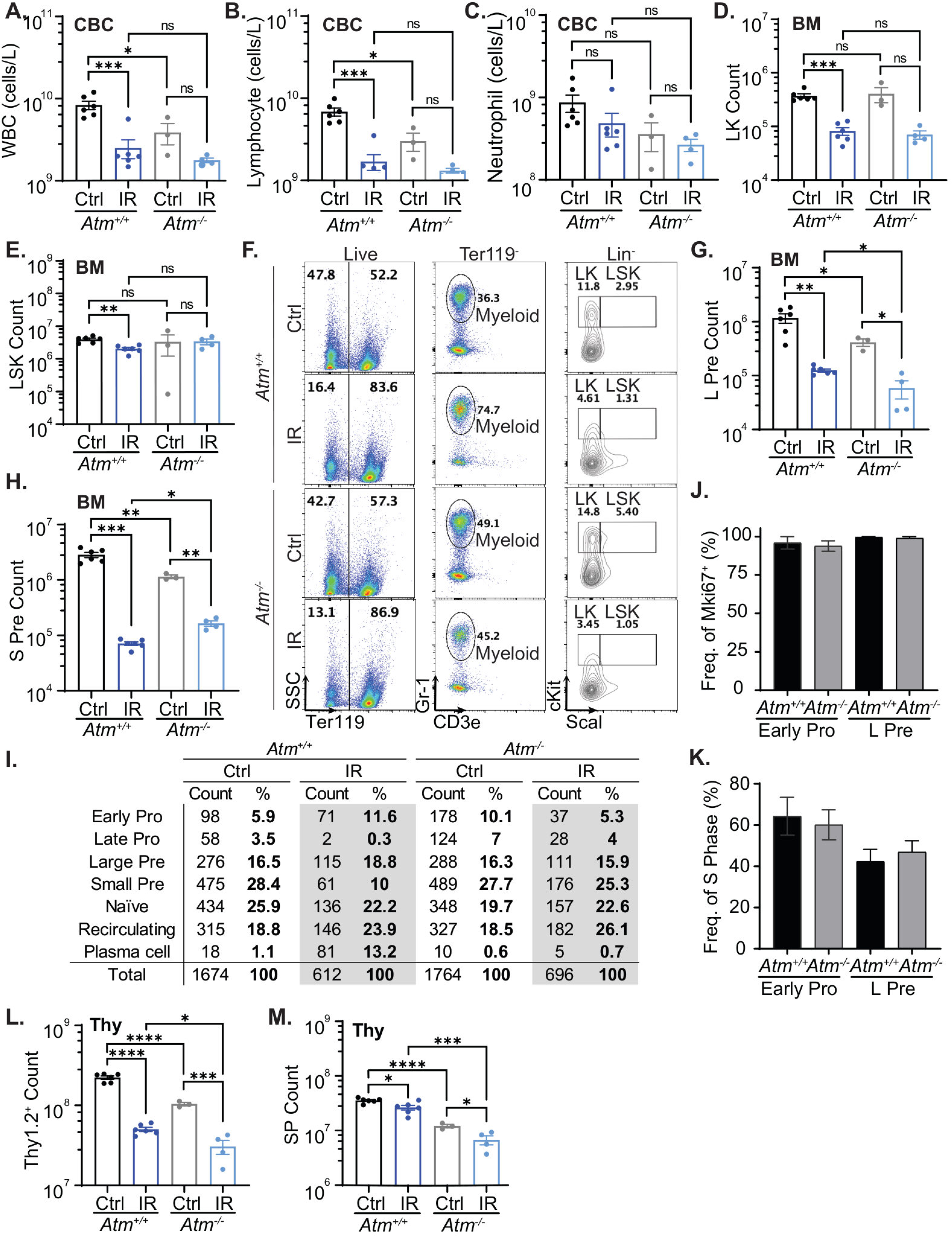
Baseline hematopoietic parameters and supporting flow cytometry and scRNA-seq analyses in Atm+/+ and Atm-/- mice. **(A-C)** Peripheral blood WBC, lymphocyte, and neutrophil counts in control and irradiated 5-week *Atm^+/+^* and *Atm^−/−^* mice. **(D-E)** Absolute LK and LSK bone marrow progenitor counts. **(F)** Representative bone marrow flow cytometry gating of live, Ter119-, myeloid, LK, and LSK populations. **(G-H)** Absolute large pre-B and small pre-B cell counts from bone marrow by flow cytometry. **(I)** Replicate-level relative frequencies and corresponding cell counts for annotated major hematopoietic populations. **(J-K)** Baseline control proliferative status of early pro-B and large pre-B cells in *Atm^+/+^* and *Atm^−/−^*mice, shown as *Mki67^+^* frequency and S-phase frequency. **(L-M)** Absolute Thy1.2^+^ thymocyte counts and SP thymocyte counts in control and irradiated *Atm^+/+^* and *Atm^−/−^* mice. Data are shown as mean ± SEM. Statistical significance was determined by unpaired two-tailed Welch’s t-test. ns, not significant; *p < 0.05; **p < 0.01; ***p < 0.001; ****p < 0.0001.

**Supplemental Table 1:** List of antibodies used for the study.

**Supplemental Table 2:** Markers Used for Single Cell RNA-seq analyses

